# Delineating the genetic regulation of cannabinoid biosynthesis during female flower development in *Cannabis sativa*

**DOI:** 10.1101/2021.11.05.464876

**Authors:** Peter V. Apicella, Lauren B. Sands, Yi Ma, Gerald A. Berkowitz

**Affiliations:** Department of Plant Science and Landscape Architecture, Agricultural Biotechnology Laboratory, University of Connecticut, Storrs, CT 06269-4163

**Author notes:** Mydecine Innovations Group Inc, 1250 S Parker Road, Denver, CO 80231. Corresponding authors: Yi Ma,; Gerald A. Berkowitz,; Agricultural Biotechnology Laboratory, 1390 Storrs Rd. Unit 4163, University of Connecticut, Storrs, CT, 06269-4163.

**Keywords:** *Cannabis sativa*, cannabinoid biosynthesis, CBDAS, GPPS, THCAS, methyl jasmonate, prenyltransferase, transcriptional regulation

## Abstract

- Cannabinoids are predominantly produced in the glandular trichomes on cannabis female flowers. There is little known on how cannabinoid biosynthesis is regulated during female flower development. We aim to understand the rate-limiting step(s) in the cannabinoid biosynthetic pathway.
- We investigated the transcript levels of cannabinoid biosynthetic genes as well as cannabinoid contents during 7 weeks of female flower development. We demonstrated that the enzymatic steps for producing CBG, which involve genes *GPPS*, *PT* and *OAC*, could be rate limit cannabinoid biosynthesis. Our findings further suggest that cannabinoid synthases, *CBDAS* and *THCAS* in a hemp and medical marijuana variety respectively, are not critical for cannabinoid biosynthesis. The cannabinoid biosynthetic genes are generally upregulated during flower maturation, which indicate glandular trichome development.
- MeJA can potentially increase cannabinoid production. We propose that biweekly application of 100 μM MeJA staring from flower initiation would be efficacious for promoting cannabinoid biosynthesis.
- Our findings suggest that the step of CBG production could rate limit the terminal cannabinoid biosynthesis. In addition, different cannabis varieties demonstrated discrete transcriptional regulation of cannabinoid biosynthetic genes.

## INTRODUCTION

*Cannabis sativa* (cannabis or *C. sativa*) is currently garnering attention for its important chemical constituents known as cannabinoids, which have a number of pharmacological properties, including chronic pain treatment, seizure mitigation, spasticity reduction, and a host of others (Brodie & Ben-Menachem, 2018; Groce, 2018). Female plants of dioecious cannabis varieties are favored for cannabinoid production since their unfertilized flowers produce cannabinoids in the highest concentration in the capitate-stalked glandular trichomes (GSTs) of high density. Terminal cannabinoids such as cannabidiolic acid (CBDA), tetrahydrocannabinolic acid (THCA), and cannabichromenic acid (CBCA) are the products of a biosynthesis pathway (Fig. S1). Cannabinoid biosynthetic enzymes (synthases for THCA, CBDA, and CBCA are generated from mRNA in the rosette of gland cells at the top of GSTs and secreted into the extracellular cavity at the top of the trichome where terminal cannabinoid synthesis and accumulation takes place (Balcke *et al*., 2017; Rodziewicz *et al*., 2019).

Although there are published studies on cannabis horticulture, trichome cell biology and the chemistry of its valuable metabolites, little is known on the regulation of cannabinoid biosynthesis in *C. sativa*. We aimed to elucidate (a) how cannabinoid biosynthesis is regulated, (b) which of the enzymatic step(s) in the biosynthetic pathway rate-limit the production of THCA and CBDA, and (c) what factors and developmental programming that occur in the flowers of female plants that could be manipulated to enhance and improve the quality of production and/or prevent occurrences such as unacceptably high THCA generation in hemp varieties cultivated for CBD production. Our experimental work attempted to bring new information and understanding to these questions and aimed to add greater resolution by measuring cannabinoid content and expression of key biosynthesis genes as the female flower matures and generates cannabinoids. Another factor that may be important for cannabinoid production is trichome density. Studies have shown that the exogenous application of the plant hormone, jasmonic acid, increases trichome density and secondary metabolite concentrations in plants containing glandular trichomes (Boughton *et al*., 2005; Yan *et al*., 2017). However, little research has been done to evaluate if applications of methyl jasmonate (MeJA) increases cannabinoid content in female flowers of cannabis plants.

In this report, we monitored transcripts of cannabinoid biosynthetic genes during 7 weeks of female flower development in two cannabis varieties, a hemp variety Cherry Wine (CW) and a medical marijuana variety Gorilla Glue (GG). Meanwhile, we measured cannabinoid concentrations at the same time points. In general, cannabinoid biosynthesis showed correlation with the expression of the corresponding gene except for THC and *THCAS* in GG. In addition, we examined the effect of MeJA on cannabinoid production. The results showed that MeJA increased THC content by about 21% percent compared to control. The rate limiting step(s) of cannabinoid biosynthesis has not been heretofore examined; results presented here address this issue.

## MATERIALS AND METHODS

### Plant materials

Cuttings of 5-month old GG mother plants were taken and rooted in an EZ-Cloner Classic ChamberTM (Sacramento, California) under aeroponic conditions. Rooted cuttings were grown under 18 h light/ 6 h darkness for 9 weeks in the closed-loop commercial facility, CTPharma LLC. Plants were then grown under 12 h light/12 h darkness for reproductive growth.

Cuttings of 5-month old CW mother plants grown in the greenhouse at University of Connecticut were taken to produce new clones for experiments. Plants were grown under 18 h light/6 h darkness for vegetative growth. After 3 weeks of vegetative growth, plants shifted to a new cycle of 12 h light/12 h darkness to initiate floral development. The plants were maintained in this environment for 7 weeks.

### Sample collection, RNA isolation, and cDNA synthesis

Approximately 0.1 g of flower material was collected and then immediately placed into liquid nitrogen for flash freezing. Samples were then stored in a −80 ° C freezer until RNA extraction. Macherey-Nagel Plant and Fungi Mini kit for RNA (Düren, Germany) was used for RNA extraction according to the manufacturer’s protocol with modifications. Single-stranded cDNA was synthesized from 1 μg of the isolated RNA using the Bio-Rad iScript™ cDNA synthesis kit (Hercules, California).

### Cannabinoid Extraction and HPLC Analysis

Approximately 0.5 g of flower material was added to individual 50 mL Falcon^®^ tubes, then 20 mL of 25°C 9:1 methanol:dichloromethane (v/v) was added to tubes. Each tube was vortexed vigorously for 3 sec, and then tubes were placed on a shaker plate for 90 min. Tubes were then vortexed vigorously again for 3 sec before transferring 1-2 mL of extract into a syringe and pressing the liquid through 0.45 μm filter into a 2 mL microcentrifuge tube. Following filtration, 100 μL of extract was added to 900 μL of methanol in autosampler vials. The extracts were stored at −20 °C until further analysis.

Cannabinoids were resolved on a 1260 Agilent High Pressure Liquid Chromatograph (HPLC) instrument (Santa Clara, California) using the Restek Raptor ARC C18 column (150 x 4.6 mm, 2.7 μm particle size) (Bellefonte, Pennsylvania) and a binary isocratic gradient mobile phase A (water, 5 mM ammonium formate, 0.1% (v/v) formic acid) and mobile phase B (acetonitrile, 0.1%(v/v) formic acid) at 30°C. A UV detector at 228 nm was implemented for the analysis. Peaks on the chromatograms were identified through comparison of retention times of cannabinoid reference RestekTM (Bellefonte, Pennsylvania) standards. Cannabinoid quantification was performed by comparing values to a calibration curve created with cannabinoid reference standards. The cannabinoid content (CBG, CBD and THC) represents the total amount of carboxylated (CBGA, CBDA and THCA) and decarboxylated cannabinoids, which is presented as the percentage of total flower dry weight.

### Real Time PCR

Reactions were performed with the Bio-Rad iTaq Universal SYBR Green Supermix (Hercules, California). Primers were chosen based off what other authors have used in other publications (Table S1). Ubiquitin (UBQ) was used as a reference gene. RT-qPCR analysis was performed using four biological replicates.

### Methyl Jasmonate (MeJA) treatment

Different concentrations of MeJA (Food Grade, Sigma Aldrich) were prepared from a stock 95%MeJA solution: 100 μM, and 500 μM, 1000 μM, and a water control. In CTPharma LLC., MeJA was applied to three biological replicates of ‘White Tangy Haze’ for each concentration to the point of runoff after they had developed flowers for 2 weeks. Each plant received approximately 1 L of MeJA solution. Weekly samples for cannabinoid analysis were harvested for the next 4 weeks.

## RESULTS

### The correlation between cannabinoid production and the transcript levels of corresponding biosynthetic genes

To learn the regulation of cannabinoid production during flowering development, we examined flowers over the course of the 7-week flower development period (Fig. S2). Week 1 is right after the transition to reproductive growth and week 7 flowers are fully mature. Cannabigerol (CBG) levels typically did not rise above 2% in any variety that we studied. In CW, CBG levels increased consistently and peaked in week 4. After week 4, CBG was maintained at a similar level (Fig. 1a, c, black line). In GG, CBG levels also peaked in week 4 but decreased thereafter over the next three weeks (Fig. 1b, d, black line). The enzyme, PT, catalyzes the alkylation of GPP and OA to create the precursor cannabinoid, CBGA. PT1 was initially found to catalyze CBGA biosynthesis (Page & Boubakir, 2012). Most recently PT4 was reported to have higher enzymatic activity than PT1 (Blatt-Janmaat & Qu, 2021). Both genes are enriched in glandular trichomes and localized in the chloroplast (Page & Boubakir, 2012; Gülck *et al*., 2020). We examined the expression of both *PT1* and *PT4* to learn how these two genes are transcriptionally regulated during female flower development.

**Figure 1.**
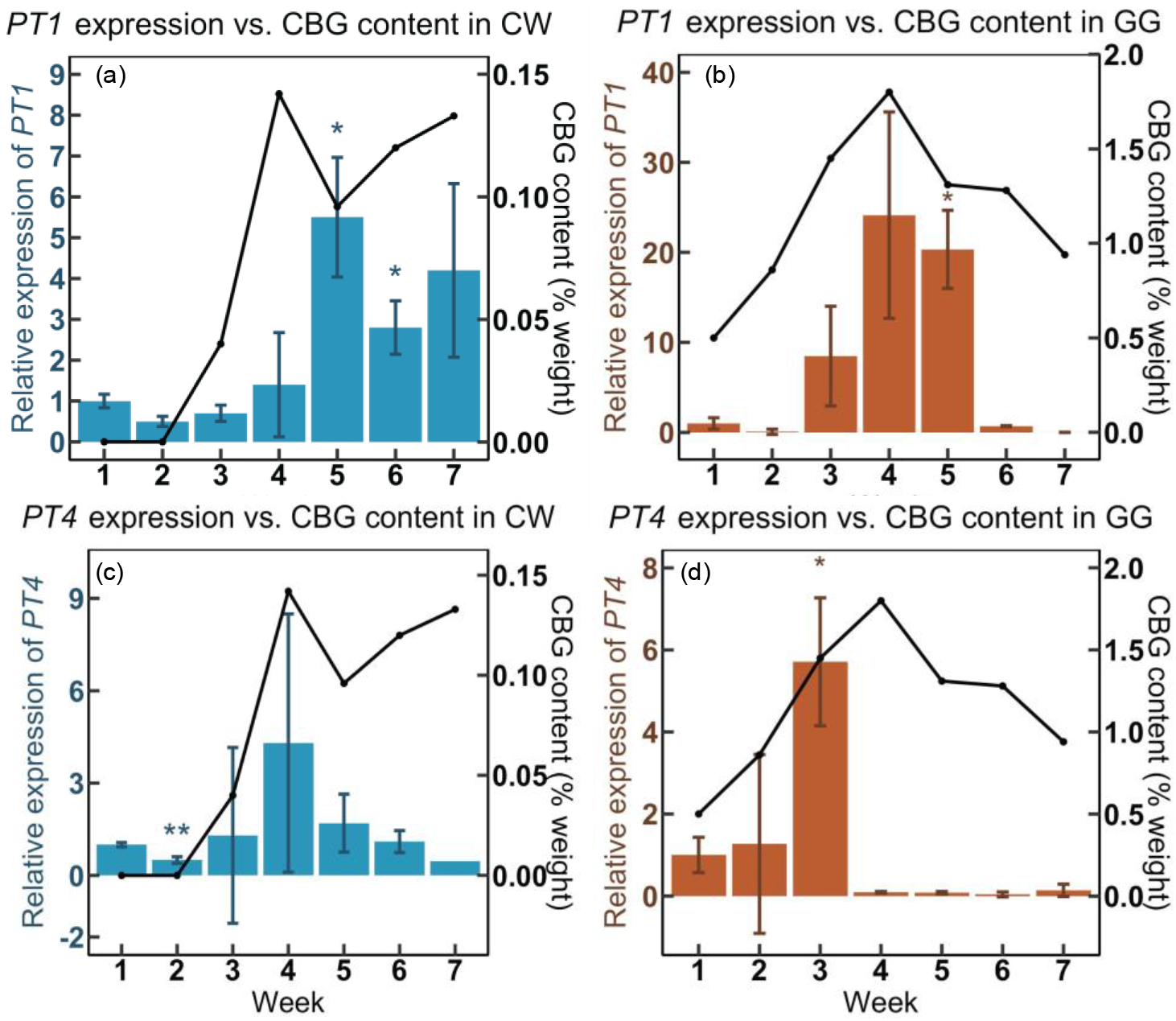
The remained CBG amount and gene expression of *PT1* and *PT4* over the course of seven-week flower development in CW (a, c) and GG (b, d). y axes on the left show relative expression of *PT1* and *PT4*. y axes on the right show the percentage of CBG amount in dry flower samples. Data are presented as means ± SE (n = 4). Significant difference of *P* < 0.05 (*) and *P* < 0.01 (**) was determined by Student *t*-test.

For both hemp (CW) and marijuana (GG) varieties, there was a general trend of increased expression of *PT1* and *PT4* (bars in Fig. 1) as the flower matured and CBG levels rose (black lines in Fig. 1). Maximal CBG levels occurred at week 4, and expression of the *PT* genes generating CBG increased during flower development (peaking at weeks 3-5). In GG flowers, expression of both *PT1* and *PT4* declined from peak levels as the flowers further matured. This trend was less pronounced in CW. Our results suggest that *PT1* expression is more correlated with CBG content.

We next examined cannabinoids and their corresponding synthase genes. CBD accumulation was steadily increased from week 1 to week 7 in CW (Fig. 2a, black line). *CBDAS* showed a similar expression pattern to that of *PT1* in CW, with the increased expression in weeks 5 to 7 being plateaued. In the marijuana variety GG, no CBD was detected despite that *CBDAS* transcripts could still be measured (Fig. 2b). Unlike the upregulation of *CBDAS* in CW, *CBDAS* expression in GG was maintained at similar levels during the whole period of flower development. *CBDAS* transcripts were detected in GG; CBDAS enzyme should be produced in GG. One possibility for the non-detected CBD is that there are mutations in the *CBDAS* gene that result in a nonfunctional protein. It is also possible that the amount of CBDAS enzyme is very low and the synthesized CBD is under the detectable level.

**Figure 2.**
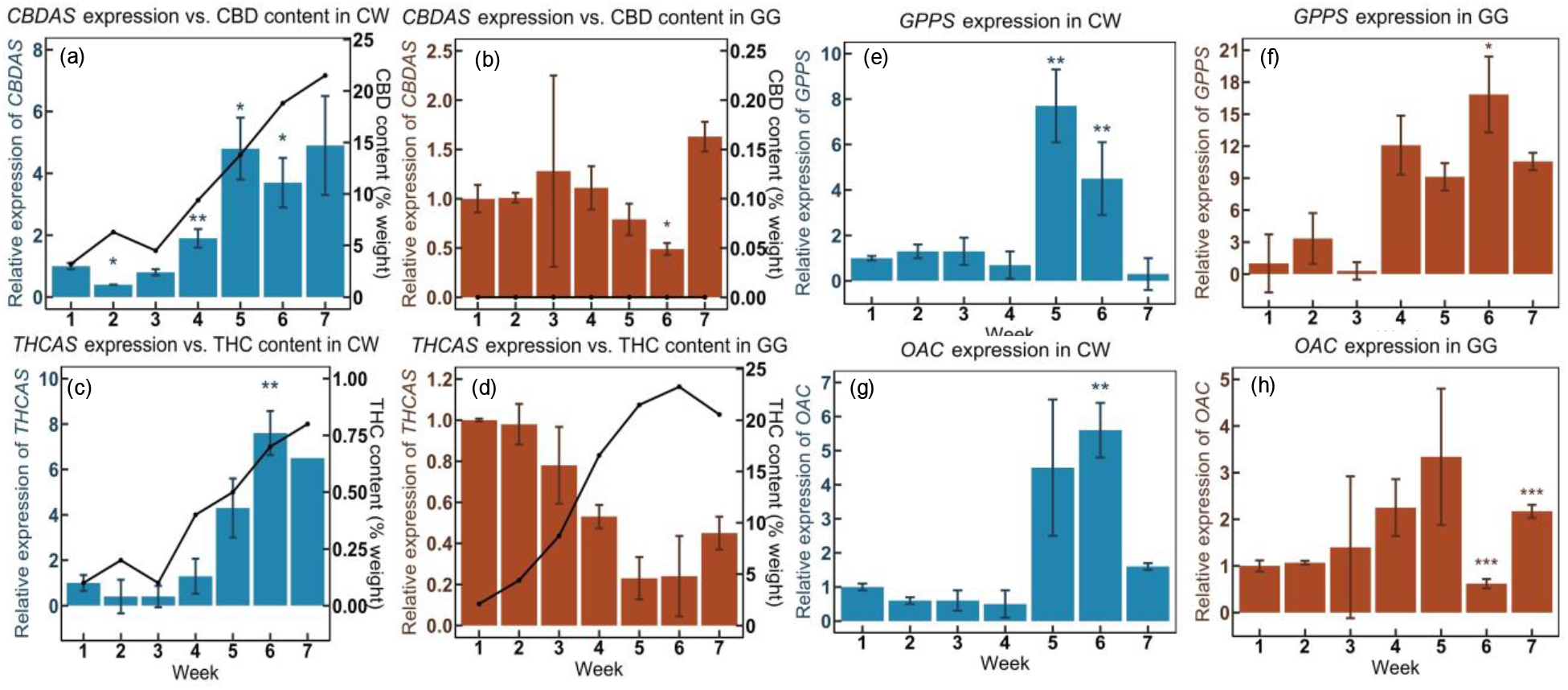
Analysis of CBD and THC accumulation with the expression of genes in the cannabinoid biosynthetic pathway over the course of seven-week flower development. (a, b) Superimposition of CBD content with *CBDAS* expression in CW (a) and GG (b). (c, d) Superimposition of THC content with *THCAS* expression in CW (c) and GG (d). Gene expression of *GPPS* and *OAC* in CW (c, g) and GG (d, h). Data are presented as means ± SE (n = 4). Significant difference of *P* < 0.05 (*), *P* < 0.01 (**) and *P* < 0.001 (***) was determined by Student *t*-test.

In CW, THC biosynthesis and *THCAS* gene expression showed a very similar trend to that of CBD production and *CBDAS* gene expression respectively (Fig. 2a). *THCAS* gene expression was significantly increased through week 4-7 in CW (bars in Fig. 2c), while it had a descending trend during flower development in GG (Fig. 2d). THC content in both CW and GG increased steadily (black lines in Figs. 2c,d). The findings on THC production indicate that THC biosynthesis does not rely on the upregulation of *THCAS* gene expression. *THCAS* transcripts may have been at higher levels at the beginning of flower development in GG.

Even though CBC possesses very low amount compared to the other two cannabinoids, *CBCAS* was still highly upregulated in both varieties (Fig. S3). CBC content did not rise above 0.5% dry weight in CW; there was no apparent increase in the rate of CBC production (data not shown).

### Regulation of upstream genes in cannabinoid biosynthetic pathway

GPPS is an enzyme that produces geranyl pyrophosphate (GPP) (Fig. S1). In CW, *GPPS* showed a similar expression pattern to the synthase genes despite that it was abolished in week 7 (Fig. 2e). *OAC* expression showed a similar pattern to that of *GPPS* in CW (Fig. 2g). GPP and OA together produce CBG, therefore it is reasonable that the two genes are similarly regulated. In GG, *GPPS* expression was highly upregulated from week 4 through week 7 (Fig. 2f). *OAC* expression did not increase as much as that of *GPPS* but still demonstrated a similar expression pattern (Fig. 2h). It is possible that *OAC* transcripts are already abundant in GG to produce sufficient OA for cannabinoid production. Because GPP is also the precursor for monoterpenes (Allen *et al*., 2019), greater upregulation of *GPPS* is still necessary to produce ample amount of GPP. The different expression patterns of *GPPS* in CW and GG also suggest discrete regulation of monoterpene production in the two varieties.

### Correlation analysis of cannabinoid biosynthetic genes and cannabinoid production

Overall, the expression of cannabinoid biosynthesis genes follows a similar trend during flower development. The genes were upregulated in the middle stages of flower development, and cannabinoid accumulation increased steadily as flowers develop. We analyzed the correlation between the expression of cannabinoid biosynthetic genes and cannabinoid content using Pearson’s correlation matrix.

In CW, it is clearly shown that the expression of *PT1* (*AP* in the figure) was positively correlated with CBG content over time. There was a statistically significant positive correlation between *PT1* expression and THC or CBD production. In addition, expression of both *CBDAS* and *THCAS* was positively correlated with CBD and THC production respectively in CW (Fig. 3a). *GPPS* has positive but weak correlation with cannabinoid production in CW. Interestingly, in GG, there was a strong negative correlation between the expression of *THCAS* and THC (Fig. 3b), suggesting that increasing *THCAS* transcript levels would not facilitate THC production. However, *GPPS* is highly correlated with THC production in GG, while the positive correlation was much weaker in CW. THC production has positive correlation with the steps for CBG production but not with *THCAS*, suggesting that CBG production is critical for THC biosynthesis, which is not dependent on *THCAS* transcript level. OA is a special component in cannabinoid biosynthetic pathway, and there is a positive correlation between *OAC* expression level and cannabinoid production in both the two varieties.

**Figure 3.**
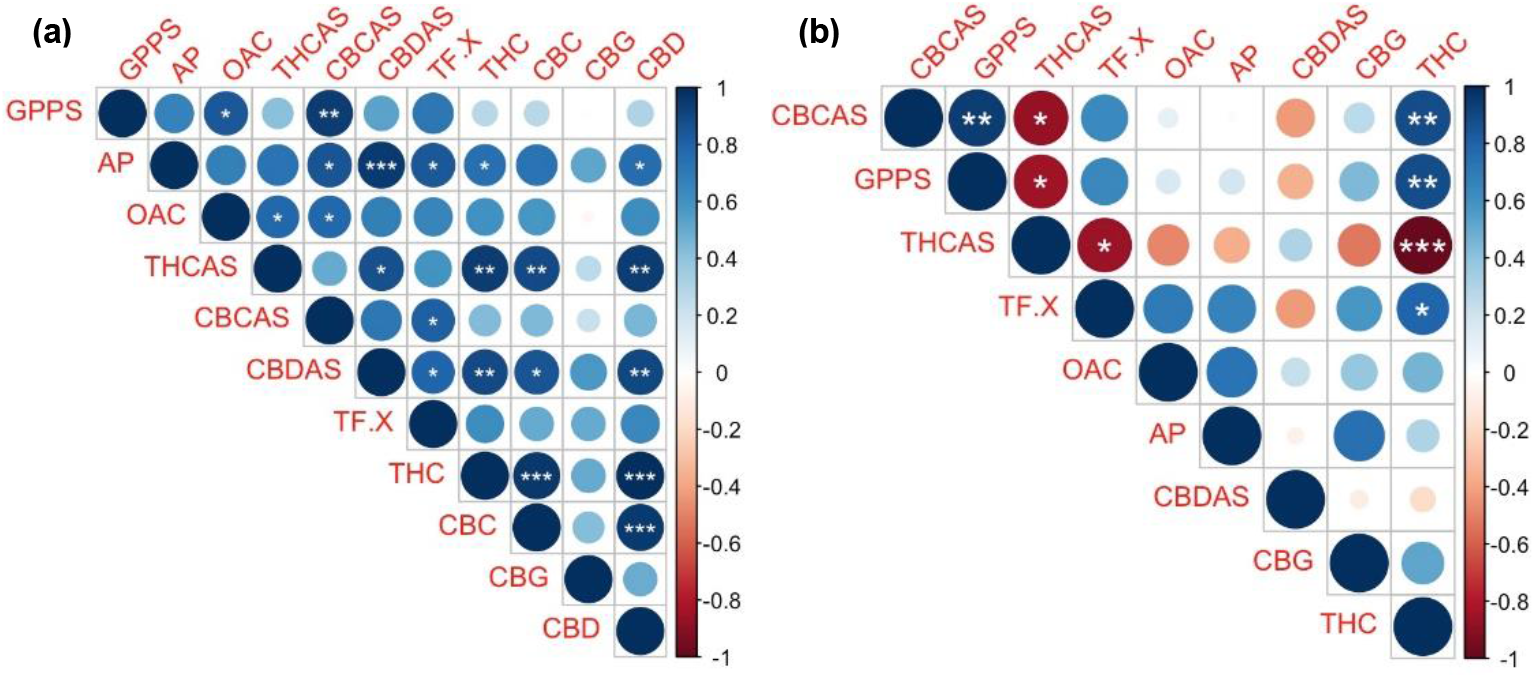
Graphic representation of correlation matrices demonstrates the correlation plot of cannabinoid biosynthetic genes and metabolites for CW (a) and GG (b). Blue indicates a positive correlation, while red indicates a negative correlation. Size and color of circle represent strength of correlation. Figures were generated using the R corrplot package https://github.com/taiyun/corrplot). (*, *P* < 0.05; **, *P* < 0.01; ***, *P* < 0.001).

### Effects of MeJA on cannabinoid production

MeJA has been shown to enhance secondary metabolites production in various plant species (Choi *et al*., 2005; Kim *et al*., 2006; Chen *et al*., 2006; Yan *et al*., 2017; Jiang *et al*., 2017). We examined how MeJA treatment could regulate cannabinoid production in a marijuana variety, ‘White Tangy Haze’.

In the first week following application, only plants sprayed with 1 mM MeJA had significantly higher THC (Fig. 4a) and total cannabinoid levels (Figure S4a) than those treated with water. All the treatments in the second week ameliorated THC production compared to the water control. More interestingly, plants treated with 100 μM MeJA produced a similarly high level of THC (Fig. 4b) and total cannabinoid contents (Fig. S4b) to those treated with 1 μM MeJA. The effect of MeJA appeared to be subsiding three weeks after application but 1 mM MeJA treatment still showed significantly higher THC production (Fig. 4c). The positive effects of MeJA treatments (100 μM and 1 mM) on cannabinoid production was compromised 4 weeks after treatment (Fig. 4d). The observation indicates that MeJA is efficacious for enhancing cannabinoid production and could be used to increase cannabinoid production in cannabis growth industry.

**Figure 4.**
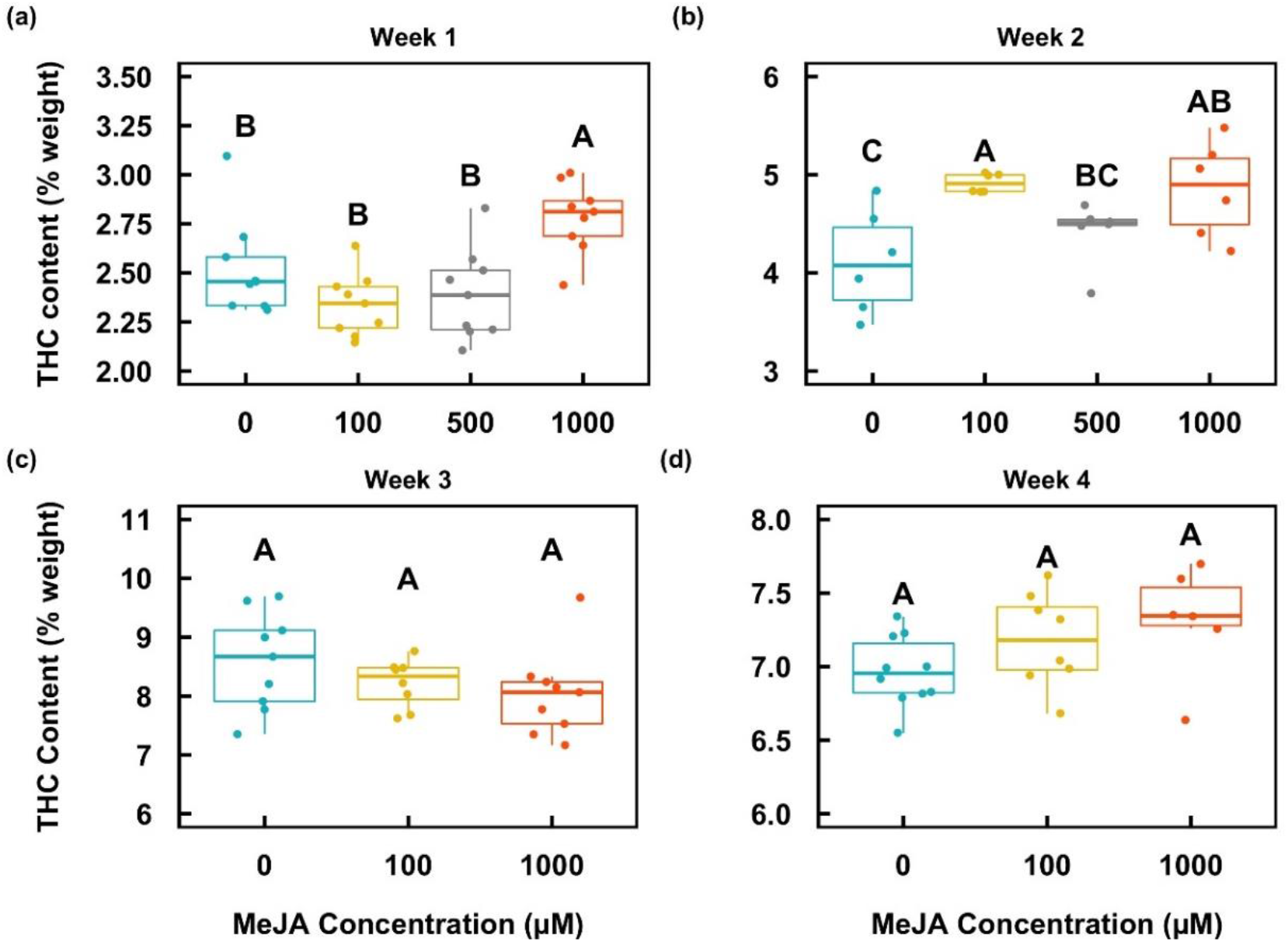
THC content in White Tangy Haze flowers over the period of four weeks (a-d) following application of MeJA. The box whiskers indicate variability outside the upper and lower quartiles (the upper and lower lines of the box) (n = 12). Different letters indicate statistical significance (*P* < 0.05) determined by one-way ANOVA followed by Fisher’s LSD.

## DISCUSSION

### CBGA biosynthesis is the rate-liming step in the cannabinoid biosynthetic pathway

We provided a comprehensive analysis on the correlation between cannabinoid production and expression of the biosynthetic genes. In general, both hemp CW and medical marijuana GG showed gene upregulation during female flower maturation, indicating that the regulation of cannabinoid biosynthetic genes is also associated with trichome development. Livingston *et al*.(2020) showed that cannabinoids are primarily accumulated in the GSTs, which are abundant during the later stages of flower development. Our gene expression and cannabinoid quantification analysis are consistent with their observation.

Our findings on *THCAS* gene expression and THC production is consistent with previous reports that *THCAS* was downregulated with the development of flowers despite the increase of THC production (Richins *et al*., 2018; Liu *et al*., 2021), which could be due to the upregulation of two transcription factors (TF) that suppresses *THCAS* (Liu *et al*., 2021). In addition, Muntendam *et al*. (2012) did not find strong correlations between synthase enzymes and their corresponding products. Onofri *et al*. (2015) demonstrated that the expression levels of *CBDAS* or *THCAS* are not associated with cannabinoid production when comparing various cannabis strains together. Our and other studies imply that the transcript level of the synthase genes is not critical for cannabinoid production. This is possibly because the varieties with high cannabinoids are already bred to have higher synthase transcript levels. Therefore, increasing the synthases alone may enhance cannabinoid production.

We further showed that *PT*, *GPPS* and *OAC* expression were also significantly increased (Figs. 1, 2e-h). Based on our findings in GG, we propose that THCA production is mainly determined by the expression and activity of *PT* and *GPPS*, the steps leading to CBG production. It has been shown in several plant species that overexpression or upregulation of GPPS increased monoterpenes (Orlova *et al*., 2009; Xi *et al*., 2016; Yin *et al*., 2017; Chuang *et al*., 2018a,b). Pearson’s matrix further demonstrated that *GPPS* is highly correlated with THCA in GG (Fig. 3b). It is reasonable to propose that promoting GPP production could enhance CBG amount and eventually cannabinoid production in cannabis. In addition, due to the dual function of GPP as a precursor, increasing *PT* expression would allow more GPP to be allocated for CBG production.

We included a TF involved in glandular trichome initiation in this report (Fig. 3). Its expression showed positive correlation with cannabinoid contents, suggesting that glandular trichome development could also be essential for cannabinoid production. The genes discussed above could be targets for traditional breeding or genetic modification.

### Hemp and marijuana differ in transcriptional regulation of cannabinoid biosynthetic genes

We examined transcript levels of cannabinoid biosynthetic genes in two types of cannabis varieties, hemp CW and medical marijuana GG. Our data suggest different transcriptional regulation of cannabinoid biosynthesis in the two varieties. *PT1* expression in CW increased in week 5 and was maintained at a similar level (Fig. 1a). *PT1* expression in GG gradually increased until weeks 4 and 5 but was completely abolished in later weeks (Fig. 1b). The findings suggest that the *PT1* genes may be distinctly regulated in hemp and marijuana. In addition, *GPPS* expression in GG and CW also showed temporally different regulation (Fig. 2e,f). Distinct monoterpene profiling could also be a feedback regulation of *GPPS* expression in the two varieties.

We showed that transcription of *THCAS* in CW is normal, but CW does not produce as much of THC as GG. To learn if there are mutations in *THCAS*, we sequenced CW *THCAS*. The result showed that CW THCAS is 100% identical to the known sequence on NCBI and fully functional. By comparing *THCAS* transcripts in CW and GG, we found that GG *THCAS* expression is hundreds of folds higher than CW *THCAS* (Fig. S5), indicating that *THCAS* is tremendously suppressed in CW. *THCAS* in CW was still upregulated and higher than illegal level of THC produced. Therefore, suppressing of *THCAS* in CW would possibly reduce THC to the legal level.

### MeJA regulates cannabinoid biosynthesis

MeJA is the plant hormone that plays important roles in secondary metabolite production and glandular trichome development (Chen *et al*., 2017; Boughton *et al*., 2005; Wang *et al*., 2010; Maes *et al*., 2011; Montiel *et al*., 2011; Yu *et al*., 2012; Shen *et al*., 2016; Yan *et al*., 2017; Hua *et al*., 2021). However, there is no publication regarding the effect of MeJA on cannabinoid production. We showed that 100 μM MeJA was as effective as 1 mM MeJA to enhance THC production two weeks after treatment (Fig. 4b). Jiang *et al*. (2017) demonstrated that high MeJA concentrations caused necrosis on cucumber leaves and did not improve monoterpene production. Therefore, for the improvement of cannabinoid production, a biweekly 100 μM MeJA treatment during flower development would be efficient to promote cannabinoid production.

Studies have shown that MeJA controls secondary metabolite biosynthesis by modulating TFs that regulate glandular trichome formation and biosynthetic genes. MeJA upregulated TFs that induce glandular trichome formation and increased artemisinin content in *Artemisia* and terpene production in tomato (Yan *et al*., 2017; Chalvin *et al*., 2020; Schuurink & Tissier, 2020). Furthermore, JA responsive TFs in various plant species directly or indirectly regulated transcription of terpene biosynthetic genes (Van Der Fits & Memelink, 2000; Li *et al*., 2015; Jiang *et al*., 2016; Shen *et al*., 2016; Nolan *et al*., 2017; Chuang *et al*., 2018a,b; Hua *et al*., 2021). The alteration of these MeJA upregulated TFs resulted in changes in secondary metabolite levels. Based on our findings, MeJA could also play vital roles in cannabinoid and secondary metabolite biosynthesis in cannabis. It would be critical to identify TFs in cannabis that are responsible for glandular trichome development and transcriptional regulation of cannabinoid biosynthetic genes.

## ACKNOWLEDGMENTS

We thank Frederick Pettit and Shelly Durocher for their assistant in maintaining cannabis plants in the greenhouse. We thank CTPharma LLC for providing cannabis plant materials for our experimental analysis. The work was supported by CTPharma LLC.

## AUTHOR CONTRIBUTIONS

PVA, YM and GAB designed the research; PVA and LBS performed experiments and analyzed data; PVA, YM and GAB wrote the manuscript; GAB and YM supervised the research.

The authors declare no conflict of interest.

## Supplemental Figure legends

Figure S1. Cannabinoid biosynthetic pathway. Purple inscriptions refer to the enzymes, including geranyl pyrophosphate synthase (GPPS), olivetolic acid cyclase (OAC), aromatic prenyltransferase (AP), tetrahydrocannabinolic acid synthase (THCAS), cannabidiolic acid synthase (CBDAS), and cannabichromenic synthase (CBCAS), that catalyze reactions. Green labels indicate the metabolites. A non-enzymatic conversion, which is expedited in the presence of heat, spurs the generation the decarboxylated cannabinoids: THC, CBD, and CBC. The figure is created using Chemdraw.

Figure S2. Representative pictures show flower development over a period of 7 weeks. Scale bar is 2 cm. Note that there is barely flower tissue developed one week after transition to short day condition.

Figure S3. Real-time PCR analysis of *CBCAS* in two C. sativa varieties, CW (a) and GG (b). Data are presented as means ± SE (n = 4). ** indicates *P* < 0.01.

Figure S4. Total cannabinoid content in White Tangy Haze flowers over the period of four weeks (a-d) following application of MeJA. The box whiskers indicate variability outside the upper and lower quartiles (the upper and lower lines of the box) (n = 12). Different letters indicate statistical significance (*P* < 0.05) determined by one-way ANOVA followed by Fisher’s LSD.

Figure S5. The comparison of *THCAS* expression in week 1 between CW and GG. *THCAS* expression is hundreds of folds higher in the medical marijuana variety GG than in the hemp variety CW. Data are presented as means ± SE (n = 4). * indicates *P* < 0.05.

